# Ancient sedimentary DNA shows rapid post-glacial colonisation of Iceland followed by relatively stable vegetation until Landnám

**DOI:** 10.1101/2021.01.15.426816

**Authors:** Inger Greve Alsos, Youri Lammers, Sofia E. Kjellman, Marie Kristine Føreid Merkel, Emma M. Bender, Alexandra Rouillard, Egill Erlendsson, Esther Ruth Guðmundsdóttir, Ívar Örn Benediktsson, Wesley R. Farnsworth, Skafti Brynjólfsson, Guðrún Gísladóttir, Sigrún Dögg Eddudóttir, Anders Schomacker

## Abstract

Understanding patterns of colonisation is important for explaining both the distribution of single species and anticipating how ecosystems may respond to global warming. Insular flora may be especially vulnerable because oceans represent severe dispersal barriers. Here we analyse two lake sediment cores from Iceland for ancient sedimentary DNA to infer patterns of colonisation and Holocene vegetation development. Our cores from lakes Torfdalsvatn and Nykurvatn span the last *c*. 12,000 cal. yr BP and *c*. 8600 cal. yr BP, respectively. With near-centennial resolution, we identified a total of 191 plant taxa, with 152 taxa identified in the sedimentary record of Torfdalsvatn and 172 plant taxa in the sedimentary record of Nykurvatn. The terrestrial vegetation at Torfdalsvatn was first dominated by bryophytes, arctic herbs such as *Saxifraga* spp. and grasses. Around 10,100 cal. yr BP, a massive immigration of new taxa was observed, and shrubs and dwarf shrubs became common whereas aquatic macrophytes became dominant. At Nykurvatn, all dominant taxa occurred already in the earliest samples; shrubs and dwarf shrubs were more abundant at this site than at Torfdalsvatn. There was an overall steep increase both in the local and regional species pool until 8000 cal. yr BP, by which time ¾ of all taxa identified had arrived. In the period 4500-1000 cal. yr BP, a few new taxa of bryophytes, graminoids and forbs are identified. The last millennium, after human settlement of the island (Landnám), is characterised by a sudden disappearance of *Juniperus communis*, but also reappearance of some high arctic forbs and dwarf shrubs. Notable immigration during the Holocene coincides with periods of dense sea-ice cover, and we hypothesise that this may have acted as a dispersal vector. Thus, although ongoing climate change might provide a suitable habitat in Iceland for a large range of species only found in the neighbouring regions today, the reduction of sea ice may in fact limit the natural colonisation of new plant species.

## 1. Introduction

Island biota are particularly vulnerable to climate change due to dispersal limitations (Harter et al., 2015; Weigelt et al., 2016). Because of arctic amplification, the earliest and most severe impact of climate change is expected at high latitudes (Bjorkman et al., 2018; CAFF, 2013). Subsequently, the fate of the flora in this region will depend on the species’ ability to track this change. While the arctic flora is assumed to have frequent long-distance dispersal (Alsos et al., 2015, 2007), oceans represent the greatest dispersal barriers (Eidesen et al., 2013). Studies of contemporary vegetation as well as modelling simulations suggest a legacy of glaciation on patterns of taxonomic richness (Stewart et al., 2016; Svenning et al., 2015), whereas pollen records indicate either Early Holocene species saturation (Giesecke et al., 2019, 2012) or dispersal lags (Felde et al., 2018). Studies of ancient DNA may shed new light on this debate as it better detects taxa and provides a higher taxonomic resolution, thereby providing a stronger method to track past species occurrence (Clarke et al., 2020; Liu et al., 2020; Rijal et al., 2020). A recent study of ancient DNA in northern Fennoscandia suggests severe dispersal lags (Rijal et al., 2020), challenging our view of a very mobile arctic flora.

Iceland is among the most isolated islands in the North Atlantic. Glacial geological evidence and numerical modelling indicate that the last Icelandic Ice Sheet reached the shelf around Iceland during the Last Glacial Maximum (LGM) (Norðdahl and Ingólfsson, 2015; Patton et al., 2017). While the renowned symposium in Iceland in 1962 (Löve and Löve, 1963) concluded that the majority of the biota must have survived the LGM *in situ*, later studies focusing on patterns of endemism and phylogeographic concluded that most species immigrated postglacially (Alsos et al., 2015; Brochmann et al., 2003). Furthermore, the estimated rate of colonisation in Iceland (1:33-214 yrs) is far greater than the Azores (1:40,000 yrs) or Hawaii (1:20,000-250,000 yrs; Alsos et al., 2015; Schaefer, 2003; Sohmer and Gustafson, 1987). Thus, the current view is that long-distance dispersal has been frequent in the amphi-Atlantic region. However, studies of phylogeography do not allow any direct dating of events, and older surviving populations may be swamped by new immigrants.

Palaeoecological data may provide more direct evidence of species at a given time and space. There are palaeoecological studies from ∼70 profiles from ∼36 sites in Iceland, the majority of them are pollen studies of peat deposits, only covering portions of the Holocene (Geirsdóttir et al., 2020; Hallsdóttir and Caseldine, 2005). Based on a review of all these records, the first occurrence could be determined for only 52 of 430 vascular plant taxa (Alsos et al., 2016). The average arrival date was 11,050 cal. yr BP, with a range from 13,000-7000 cal. yr BP. This average arrival date coincides with a period characterised by a high concentration of sea ice from the different source regions. Further, a close link was found between sea ice and driftwood. Thus, we hypothesise that sea-ice rafting was likely an important dispersal mechanism. Sea ice may act as a bridge, which allowed the Arctic fox to migrate to Iceland, potentially carrying seeds in the fur or digestive system (Graae et al., 2004). Also, it provided a smooth surface which seeds could have been wind-carried over. Furthermore, it may act as a ferry, transporting debris, driftwood and potentially propagules attached to it (Johansen and Hytteborn, 2001; Panagiotakopulu, 2014; Savile, 1972). However, determining the first arrival date for more taxa is needed to test this hypothesis.

The oldest post-glacial palaeoecological record from Iceland is from Lake Torfdalsvatn on the Skagi peninsula. The rich pollen and macrofossil record shows a relatively diverse arctic vegetation from 13,000 cal. yr BP, with more than ten taxa surviving the Younger Dryas, indicating that these species would also be able to survive the LGM *in situ* (Rundgren, 1998, 1995; Rundgren and Ingólfsson, 1999). From around 11,300 cal. yr BP, indicators of disturbed ground like *Oxyria/Rumex* and *Koenigia* are found together with *Betula,* as well as scattered records of *Salix* and *Empetrum* (Rundgren, 1998; Rundgren and Ingólfsson, 1999). A rapid Early Holocene warming led to exposure of land and development of soils 10,500-8500 cal. yr BP (Langdon et al., 2010). The Saksunarvatn tephra (10,200 cal. yr BP) marks the transition from the Preboreal to the Boreal period, and is suggested to have temporarily reduced the vegetation diversity and pollen concentrations (Rundgren, 1998). Burial by the tephra and subsequent aeolian processes are thought to have altered vegetation communities towards species more tolerant of sand burial (Eddudóttir et al., 2015). However, pollen records indicate a recovery towards pre-deposition vegetation communities within 100 years (Caseldine et al., 2006; Eddudóttir et al., 2015; Rundgren, 1998). An increase in pollen concentrations as well as a shift from grass dominance to *Salix* occurs during the early Boreal period. This is followed by a shift to dominance of *Juniperus communis* together with *Betula nana* and *Empetrum*, until birch woodland developed from 9400-6850 cal. yr BP (Eddudóttir et al., 2015; Hallsdóttir and Caseldine, 2005; Rundgren, 1998). During the Middle to Late Holocene, woodland became more patchy or open, while heath and mires expanded from around 7000 cal. yr BP (Eddudóttir et al., 2015; Hallsdóttir and Caseldine, 2005). The Norse settlement (Landnám) from around AD 870 was a rapid process, and by AD 930, most of the farmable land was claimed (Smith, 1995; Vésteinsson and McGovern, 2012). Landnám is characterised by a rapid decline in birch pollen, especially in the lowlands (Erlendsson, 2007; Hallsdóttir, 1987; Hallsdóttir and Caseldine, 2005). This pattern is also found in the less studied highlands (Eddudóttir et al., 2020), although some northeastern sites show decrease in birch prior to Landnám (Roy et al., 2018). Following settlement, the vegetation changed due to introduced plant species, habitat diversification as a result of land-use (mainly grazing), soil erosion and increased landscape openness, and diminished competition from dominant species (mainly birch). Such environmental modifications have almost certainly led to altered local plant composition and landscape-scale distribution patterns of vegetation, and may explain increased palynological richness near farms following Landnám (Erlendsson, 2007; Möckel et al., 2017; Þórhallsdóttir, 1996).

Here we study sedimentary ancient DNA (*sed*aDNA) of two lakes to explore patterns of species arrival and vegetation development over the Holocene. We investigate the lowland Lake Torfdalsvatn, as it holds the oldest known lacustrine sedimentary record from Iceland (Axford et al., 2007; Björck et al., 1992; Rundgren, 1998, 1995; Rundgren and Ingólfsson, 1999). Furthermore, sedimentary algal pigments, stable isotopes, and biogenic silica have been analysed from the lake record (Florian, 2016). Lake Nykurvatn is located in a higher elevation catchment and represents a less studied region of Iceland. Unlike Torfdalsvatn, the site likely deglaciated during the Early Holocene (Norðdahl and Ingólfsson, 2015; Patton et al., 2017). We focus on 1) the proportion and functional type of species present prior to the onset of Early Holocene warming, which could indicate glacial survival, 2) patterns of arrival for plants of different functional groups, and 3) any modification of the vegetation related to Landnám.

### 2.1. Torfdalsvatn, North Iceland

Torfdalsvatn (66.06°N, 20.38°W, 40 m a.s.l.) is located at the Skagi peninsula in North Iceland (Fig. 1). The northern part of Skagi is characterised by gently undulating terrain with thin sediment cover. From 1961-1990, the mean annual air temperature at Hraun located on Skagi was 2.5°C, and the mean annual precipitation was 475 mm. In July, the mean air temperature at Hraun was 8.2°C (Icelandic Meteorological Office, 2020a).

**Fig. 1.**
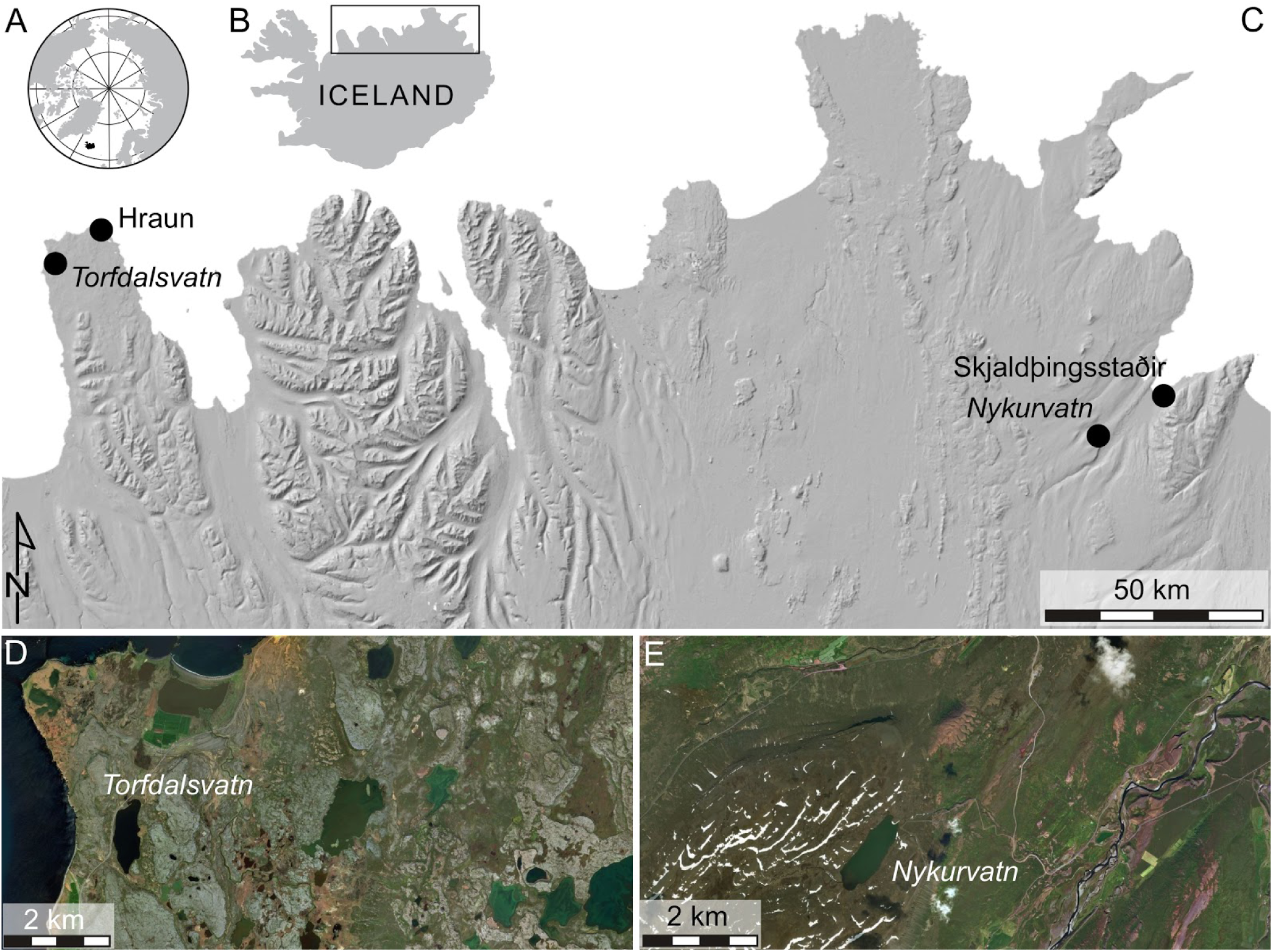
A. Overview map of the Arctic. Iceland is marked in black. B. Map of Iceland. The box indicates the extent of C. C. Terrain shaded relief map of Northeast Iceland showing the location of the two study areas. D. The area around Torfdalsvatn shown on an aerial orthophotograph. E. The area around Nykurvatn shown on an aerial orthophotograph. The imagery in C-E is available from the National Land Survey of Iceland (2020).

Torfdalsvatn has an area of 0.4 km^2^ (Fig. 2), a catchment of *c*. 3.7 km^2^ (Rundgren, 1995) and a median slope of 1.1° (Florian, 2016). The water depth at the centre of the basin is *c*. 5.1 m (Rundgren and Ingólfsson, 1999) to 5.8 m (Florian, 2016). Dwarf-shrub heaths, herb and grass tundra, and *Carex*-dominated fens characterise the vegetation in the modern catchment (Rundgren, 1997). The most abundant dwarf-shrub species are *Empetrum nigrum* and *Salix herbacea,* and the shrub *Betula nana* is also common. (Björck et al., 1992) obtained an 11.95 m long sediment sequence from Torfdalsvatn, and showed that the sedimentary record goes back to the Allerød, and that the site was ice free during the Younger Dryas. Palaeobotanical data indicate a sparse to discontinuous cover of graminoids, herbs, shrubs and dwarf shrubs during Allerød (Rundgren, 1997). During the Younger Dryas, the shrub and dwarf-shrub vegetation disappeared, and the plant cover was sparse. In the Preboreal, the graminoids, shrubs and dwarf shrubs reappeared.

**Fig. 2.**
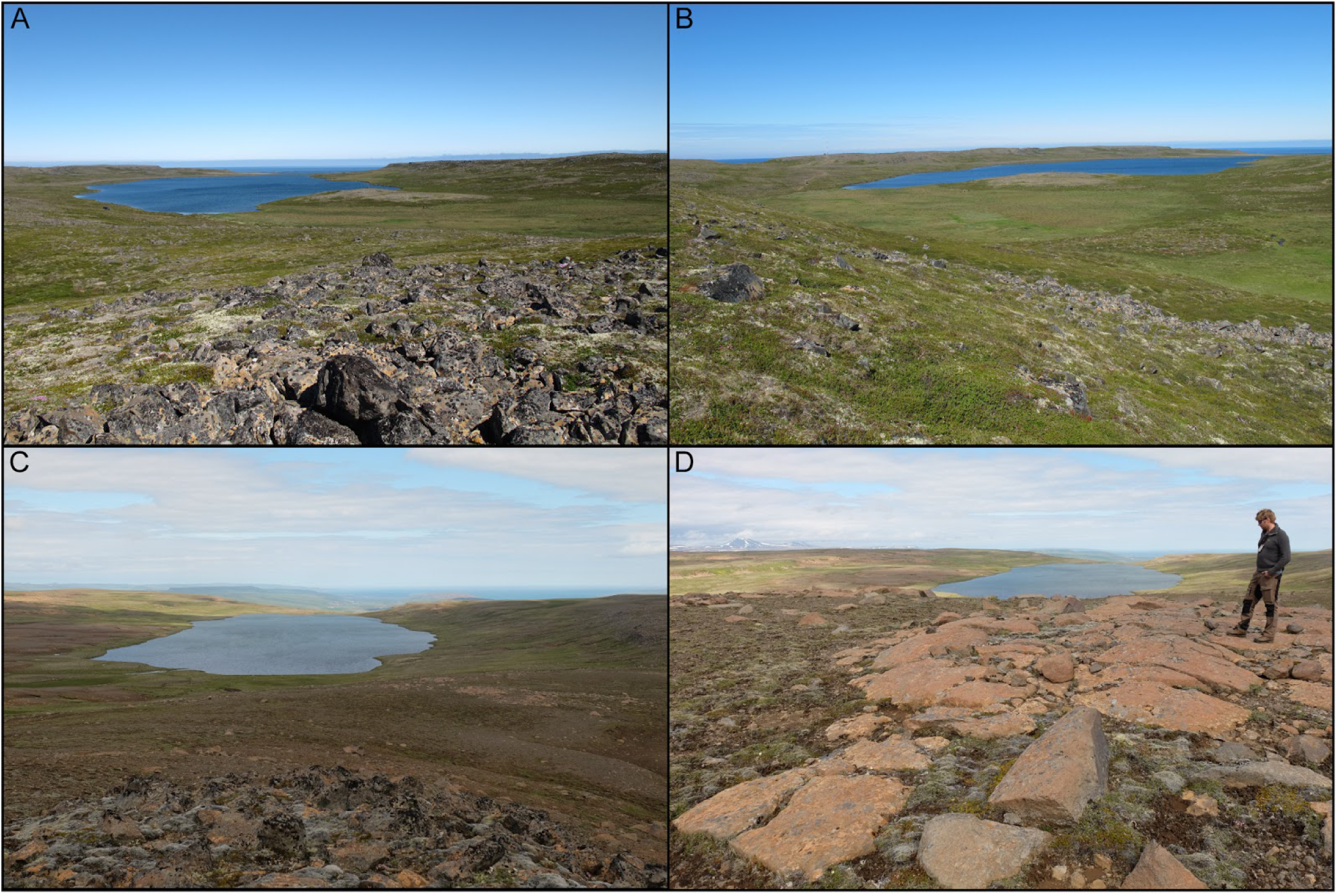
Catchment overviews of Torfdalsvatn (A-B), and Nykurvatn (C-D). Photographs taken June 2013 (G. Gísladóttir) and July 2017 (Í.Ö. Benediktsson), respectively.

Torfdalsvatn is shared between the landholdings of the active farms Tjörn and Hafnir. The farms are both featured in a letter from AD 1285 regarding land boundaries of farms owned by the Þingeyrarklaustur monastery (DI, 1857-1986, pp. 249-250). In *Jarðabók*, a land registry from the beginning of the 18^th^ century, the farms were still operating. Livestock consisted of cattle, sheep and horses. Both farms also had access to *rifhrís* (*Betula nana*) for firewood (Magnússon and Vídalín, 1927, pp. 473-479). Throughout, the inhabitants of both farms have probably operated a standard pastoral farming.

### 2.2. Nykurvatn, Northeast Iceland

Nykurvatn (65.63°N, 15.14°W, 428 m a.s.l.) is situated at the Bustarfell plateau above the Hofsárdalur valley, located in the up to 20-km wide Vopnafjörður fjord-valley system in Northeast Iceland (Fig. 1). The minimum age of deglaciation of the highland plateaus above the Hofsárdalur valley is 9400 cal. yr BP (Sæmundsson, 1995). From 1995-2019, the mean annual air temperature and precipitation at Skjaldþingsstaðir (42 m a.s.l.) in the Hofsárdalur valley, 16 km northeast of Nykurvatn, are 3.8°C and 1215 mm, respectively. The July mean temperature was 10.0°C (Icelandic Meteorological Office, 2020b). Nykurvatn has an area of 0.59 km^2^ (Fig. 2), the catchment is *c*. 10 km^2^, and the water depth at the centre of the lake is *c*. 10.0 m. The area around Nykurvatn is classified as moors and heathland (National Land Survey of Iceland, 2020).

Ownership of Nykurvatn is shared between three currently occupied farmlands, Hauksstaðir, Teigur and Bustarfell. Place names and house ruins indicate a greater density of farms and cottages in the past. The adjacent farm Hof is listed in *Landnámabók* (Book of Settlements) among the earliest settled farms (Benediktsson, 1968, p. 290) and Hauksstaðir (Haugsstaðir) and Bustarfell enter the historical records no later than in the 14th century AD. Place names and historical sources also attest to the existence of woodlands in the past (DI IV, p. 217). Given the high altitude of Nykurvatn, summer grazing of livestock probably took place by the lake.

## 3. Methods

### 3.1. Sediment coring and subsampling

Lake sediment cores were retrieved using a hand-held piston corer (150 and 200 cm long and 75 and 60 mm diameter coring tubes for Torfdalsvatn and Nykurvatn, respectively) from the lake ice. Seven core sections were collected from Torfdalsvatn (TDV2-1 to TDV2-7; water depth 5.1 m) in 2012. TDV core sections were purposely collected without overlap. Four overlapping core sections (NYK1 to NYK4; water depth 10.0 m) were collected in the centre of Nykurvatn in 2018. Additionally, one core (NYK5; water depth 3.8 m) was sampled in the distal (NE) end of the lake where the sedimentation rate is lower. The aim of this was to obtain older sediments because coring deeper below NYK4 was not possible with the piston corer. Cores were preserved in the dark and cold (∼4°C) until opening.

Half-cores were sub-sampled for *sed*aDNA immediately upon opening, wearing full bodysuits, masks and sterile gloves in dedicated clean-lab facilities with no-PCR products. The TDV cores were sampled using 5 mL sterile disposable syringes as described by Voldstad et al. (2020) whereas the NYK core was sampled using clean disposable plastic knives and spatulas as described in Rijal et al. (2020). All samples were stored at 4°C until DNA extractions.

### 3.2 Geochemical analyses

The TDV cores were logged and analysed in the sediment lab and ITRAX core facility at the University of Copenhagen (Centre for GeoGenetics) and the NYK cores at UiT The Arctic University of Norway (Department of Geosciences). ITRAX and Avaatech core scanners were used to collect high-resolution data (X-ray fluorescence, (XRF), magnetic susceptibility (MS), and optical and radiographic images) from split core sections from TDV and NYK, respectively. XRF measurements were conducted every 1 mm at 30 kV and 30 µA with a 30 s exposure for the ITRAX scanner and every 2 mm at 10 kV and 30 µA with a 20 s exposure time for the Avaatech scanner. Both core scanners have an X-ray source with a Rh (rhodium) anode (Croudace et al., 2006; Forwick, 2013). For the ITRAX measurements, Ti was normalised against the incoherent (inc) and coherent (coh) Rh scatter (Ti/(inc+coh)) to remove instrumental effects (Kylander et al., 2011). The Ti signal from the Avaatech scanner was normalised against the sum of Al, Si, S, K, Ca, Ti, Mn and Fe (Ti/Sum) to minimise artefacts (Weltje and Tjallingii, 2008). The Ti signal of the NYK core sections is presented to identify the overlap between core sections (Supplementary Fig. S1). Loss on ignition (LOI; Heiri et al., 2001) was measured to determine the total organic content for the NYK core. Samples (2 cm^3^) were collected every 5 cm, with additional samples at visible transitions in the lithology. The samples were dried at 110°C for 2 h and ignited at 550°C for 4 h.

### 3.3 Age-depth modeling and core correlation

The chronologies of both lake cores were constructed based on accelerator mass spectrometry (AMS) ^14^C measurements on plant macrofossils and identified tephra markers. The ^14^C samples were analysed at the Ångström Laboratory, Uppsala University, Sweden. For calibrating the ^14^C ages, the online OxCal software (v.4.4, Bronk Ramsey, 2009) and the IntCal20 dataset (Reimer et al., 2020) were used. All calibrated radiocarbon ages are presented as calibrated year before present (cal. yr BP; BP = 1950). Visually identified tephra layers were sampled and analysed for major element composition at the Institute of Earth Science, University of Iceland, using Electron Probe Microanalysis (EPMA). A JEOL JXA-8230 Super probe with an acceleration voltage of 15 kV, a beam current of 10 nA and a beam diameter of 10 µm was used for the EPMA. Natural and synthetic minerals were used for standardisation as well as basaltic (A99) and rhyolitic glass (Lipari Obsidian; ATHO). On each polished thin section, 20-30 point analyses were performed on randomly selected lines. The dataset was examined for outliers and contamination by microlites. All analyses with sums lower than 95% were discarded. Tephra layers were identified based on their chemical composition.

The NYK1-NYK4 core sections were stratigraphically correlated based on identified tephra marker layers, ^14^C ages, trends in the XRF data and visual similarities in lithology. Tie points were used to align the cores in AnalySeries (v. 2.0.4.2, Paillard et al., 1996) and construct a common stratigraphic depth-scale (Supplementary Fig. S1). The alignment was primarily based on the Ti/Sum ratio. We assume neither overlap nor hiatus between NYK4 and NYK5. As previously described, the TDV core sections were sampled without overlap, in a continuous sequence, and therefore do not need to be correlated.

For establishing the chronologies of the composite cores, the Bacon package (v. 2.5.0, Blaauw and Christen, 2013) with the IntCal20 dataset (Reimer et al., 2020) were used in R (v. 4.0.3, Core, 2015). The age-depth models were constructed using a mean accumulation rate (acc.mean) of 20 yr/mm for both cores and upper (d.min) and lower (d.max) depths set to the upper- and lowermost depths of the cores. For >5 cm thick tephra layers (Saksunarvatn in TDV and V1477 in NYK), a slump was added to reflect the increased sedimentation rate at deposition.

### 3.6 DNA extraction and amplification

DNA analyses were performed in the ancient DNA laboratory at the Arctic University Museum, Tromsø. Initially, 8-10 g of sediments were homogenised. DNA was extracted from subsamples of ∼0.3 g using PowerSoil Power Lyser kit and incorporating a bead beating step as in Alsos et al. (2020). We included negative controls during sampling from the core, extraction, and PCR setup, as well as positive controls during PCR. In total, 146 samples, 14 sampling and extraction negative controls, 4 PCR negative and 4 PCR positive controls were analysed.

We amplified the short and variable P6 loop region of the chloroplast trnL (UAA) intron (Taberlet et al., 2007), following the same analysis protocol Alsos et al. (2020), and running 8 PCR replicates on each DNA extract using unique 8 or 9 bp long tags added to the 5’ end of each primer. We pooled the PCR replicates, and thereafter cleaned and quantified them using Qubit (Invitrogen™ Quant-iT™ and Qubit™ dsDNA HS Assay Kit, Thermofisher). We converted the pools into DNA libraries using a Truseq DNA PCR-free low-throughout library prep kit (Illumina). The library was quantified by qPCR using the KAPA Library Quantification Kit for Illumina sequencing platforms (Roche) and a Quantstudio 3 (Life Technologies). The library was normalised to a working concentration of 10 nM using the molarity calculated from qPCR adjusted for fragment size. Sequencing was conducted on an Illumina NextSeq 500 platform (2×150 bp, mid-output mode, dual indexing) at the Genomics Support Centre Tromsø (UiT).

### 3.7 DNA sequencing analyses and filtering

We aligned, filtered and trimmed the next-generation sequencing data using the OBITools software package (Boyer et al., 2016) and a custom R-script (available at https://github.com/Y-Lammers/MergeAndFilter). The new reference library PhyloNorway, consisting of 2051 specimens of 1899 vascular plant taxa from Norway and the Polar Regions was used (Alsos et al., 2020). Additionally, we used the circumarctic/circumboreal ArcBorBryo library, containing 2445 sequences of 815 arctic and 835 boreal vascular plants as well as 455 bryophytes (Soininen et al., 2015; Sønstebø et al., 2010; Willerslev et al., 2014). Our third reference library was obtained by running the *ecotag* program on the EMBL (rl143) nucleotide reference database. We only retained sequences with 100% match to one of the reference libraries and a minimum of 3 replicates across the dataset. As 94% of the taxa in Iceland also are found in Norway (Alsos et al., 2015), match to PhyloNorway was given priority for vascular plants, whereas match to ArcBorBryo was given priority for bryophytes. For sequences assigned to several taxa, the most likely taxa with 100% match were listed based on known presence in the native flora.

We manually cross-checked that identifications were consistent among the reference libraries used, and inconsistent taxonomic assignments were cross-checked using NCBI BLAST. As this process is time-consuming, we only checked sequences that occurred with a minimum 10 replicates if they had 100% match to one of the regional reference libraries (PhyloNorway or ArcBorBryo). We followed the taxonomy of Wąsowicz (2020). Furthermore, the quality of each sample was assessed following the method described by Rijal et al. (2020).

### 3.8 Data analyses

Initial diagrams were plotted in R version 4.0.3 using the rioja (v0.9.26) and ggplot2 (v3.3.2) packages. We explored zonation for DNA using constrained incremental sum of squares (CONISS) as implemented in rioja. Final diagrams were constructed using Tilia v.2.6.1 (https://www.tiliait.com/).

## 4. Results

### 4.1 Chronology and core correlation

All radiocarbon ages, calibrated median ages and 2σ age intervals are presented in Table 1, and the tephra marker layers in Table 2. The identified tephra layers have been correlated to source volcanic system and particular volcanic event using geochemistry, stratigraphic position and age. Geochemical correlation of Hekla 1104, Hekla 3, Hekla 4 and Saksunarvatn ash in TDV and V1477, Hekla 3 and Hekla 4 in NYK are shown on elemental bi-plots in Supplementary Fig. S2. The complete geochemical dataset and standard runs are presented in Supplementary Table S1. Given the geochemical similarity of the Saksunarvatn/G10ka series tephra (Óladóttir et al., 2020) and the relatively short phase of associated volcanism compared to the Holocene scale of this study, we refer to the tephra horizon as a marker layer according to the Greenland ice core chronology (Rasmussen et al., 2006). Radiocarbon age Ua-65341 from NYK (Table 1) is an outlier when compared to Ua-67058 and tephra marker layers V1477 and Hekla 3 stratigraphically below, and was excluded when running the final age-depth model for NYK (Fig. 3B).

**Fig. 3.**
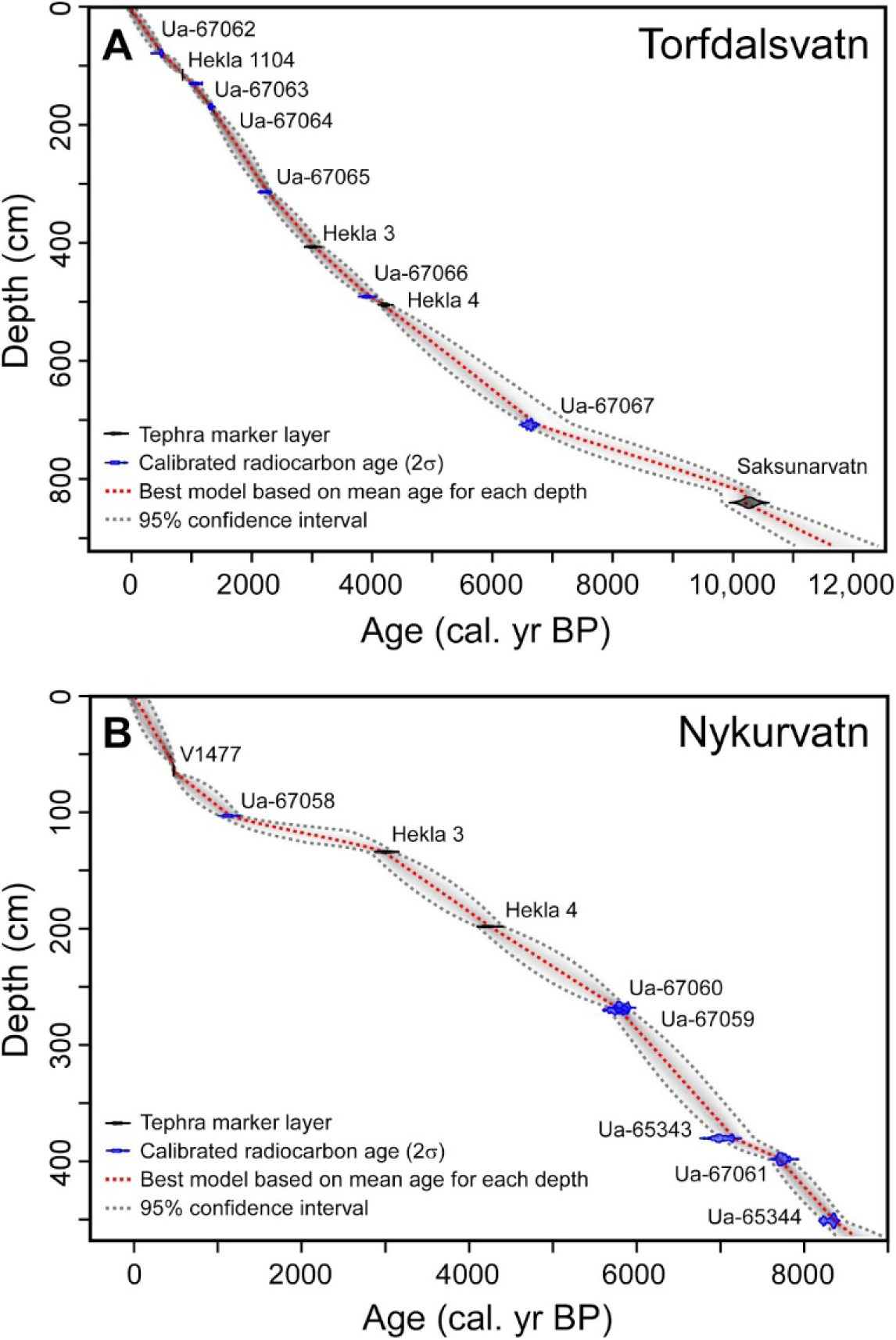
Age-depth models for Torfdalsvatn (A), and Nykurvatn (B). Details for all radiocarbon ages are given in Table 1 and tephra marker layers in Table 2 and Supplementary Table S1.

**Table 1.**
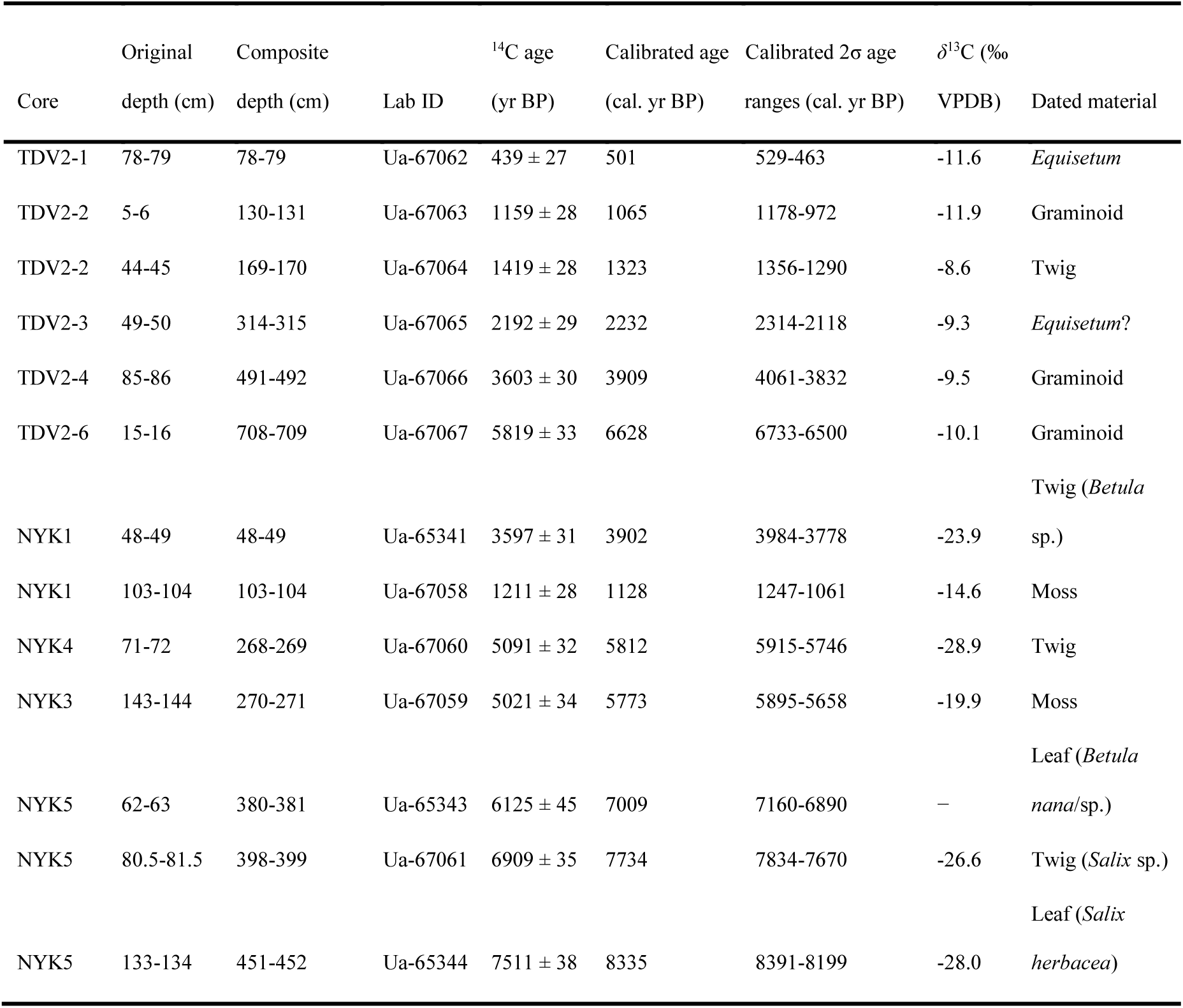
Radiocarbon ages from Torfdalsvatn (TDV) and Nykurvatn (NYK). Calibrated ages are median ages within the 2σ age ranges.

**Table 2.** Identified tephra marker layers from Torfdalsvatn (TDV) and Nykurvatn (NYK), used for the age-depth models. H = Historical; R = Radiocarbon; I = Ice core.

| Tephra layer<br>name (core,<br>composite depth) | Volcanic<br>system | Silicic (S)<br>Basaltic (B) | Tephra<br>marker | Age<br>(cal. yr BP) | Age<br>(yr BP) | Type<br>of date | Reference for age |
| --- | --- | --- | --- | --- | --- | --- | --- |
| NYK61 | Veiðivötn-Bárðarbunga | B | V1477 | 470 | - | H | (Larsen et al., 2002) |
| TDV115 | Hekla | S | Hekla 1104 | 850 | - | H | (Larsen et al., 2002) |
| TDV407,<br>NYK136 | Hekla | S | Hekla 3 | 3005 ± 57 | 2879 ± 34 | R | (Dugmore et al., 1995) |
| TDV505,<br>NYK198 | Hekla | S | Hekla 4 | 4196 ± 33 | 3826 ± 12 | R | (Dugmore et al., 1995) |
| TDV839 | Grímsvötn | B | Saksunarvatn Ash | 10267 ± 89 | - | I | (Rasmussen et al., 2006) |

The TDV sections were sampled without an overlap, and stacked on top of each other to establish the composite record. The age-depth model supports this approach. The 915 cm long TDV record spans the last *c*. 12,000 years (Fig. 3A). The correlation of the NYK core sections showed large overlaps between NYK1, NYK2 and the upper half of NYK3 (Supplementary Fig. S1). Also, part of NYK3 and NYK4 were overlapping, whereas there was no overlap between NYK4 and NYK5. Presuming no hiatus between NYK4 and NYK5 resulted in a fairly linear age-depth model for that part of the record, and we therefore consider it a reasonable assumption. The 464 cm long composite NYK record spans the last *c*. 8600 years (Fig. 3B).

### 4.2 Lithology and stratigraphy

Four main sedimentary facies were identified in Torfdalsvatn: clay, clayey gyttja, laminated gyttja, and tephra (Fig. 4A; Supplementary Figs S3-S9). The clay facies is grey and homogeneous, and the MS and Ti/(inc+coh) are high. We interpret this facies as clay formed from suspension settling of particles originating from sediment-laden streams flowing into the basin. The clayey gyttja is brown-grey and the MS and Ti/(inc+coh) records indicate a lower minerogenic content than in the clay. We interpret this facies as formed by organic sedimentation and inflow of minerogenic material by runoff from the catchment (e.g., (Björck et al., 1992; Rundgren, 1995)). The laminated gyttja facies is light olive-brown and contains numerous silt-sand sized tephra beds. The magnetic susceptibility (MS) and the Ti/(inc+coh) are very low, indicating less inflow of minerogenic material compared to the clayey gyttja. The tephra facies appear as 1-180 mm thick, black or white beds of silt-sand size volcanic glass. Angular tephra grains and characteristic chemical composition suggest that they are not reworked. Therefore, we interpret the tephra as originating from airborne material during volcanic eruptions (e.g., Gudmundsdóttir et al., 2012; Larsen and Eiríksson, 2008; Lowe, 2011).

**Fig. 4.**
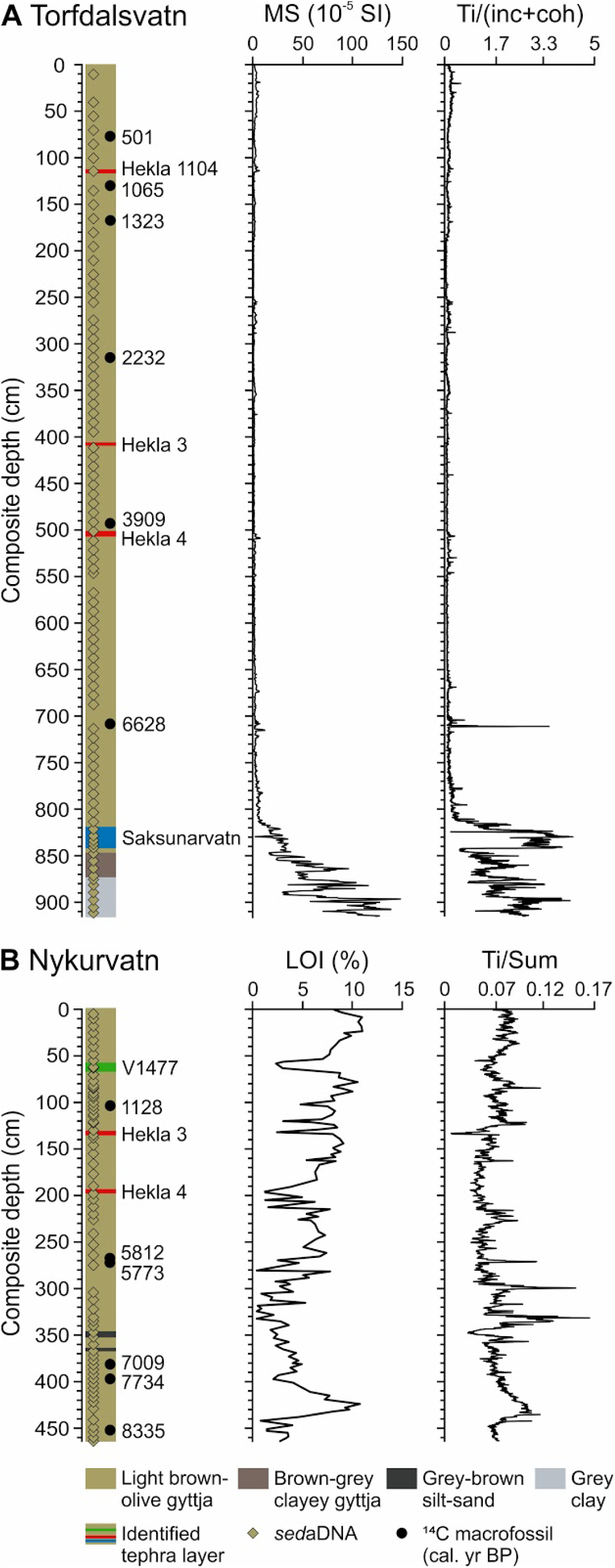
Lithology, depths for *sed*aDNA samples, identified tephra marker layers and calibrated median ages for (A) Torfdalsvatn (TDV) and (B) Nykurvatn (NYK). In addition, magnetic susceptibility (MS) and Ti/(inc+coh) are presented for TDV, and LOI (%) and Ti/Sum for NYK.

Three main sedimentary facies were identified in Nykurvatn (Fig. 4B; Supplementary Figs S10-S14). The laminated gyttja facies is light olive-brown, containing silt-sand sized tephra beds. It has an organic content of 1-11%, with the lowest values in the tephra and highest values in the massive parts of the gyttja. As in Torfdalsvatn, we interpret the gyttja facies as formed by organic sedimentation and inflow of minerogenic material by runoff. The tephra facies are 1-180 mm thick, black or white beds of silt-sand size. Again, we interpret it as originating from airborne material from volcanic eruptions (Gudmundsdóttir et al., 2012; Larsen and Eiríksson, 2008; Lowe, 2011). The grey-brown silt-sand facies appear as 3-5 cm thick beds and only occur in NYK5. We interpret this facies as deposited from higher input of minerogenic material to the basin e.g., by runoff and/or aeolian activity.

### 4.3 *sed*aDNA data

In total, 25,662,641 merged paired-end reads were obtained for both lakes. Out of these, 16,383,211 reads, belonging to 324 sequences were retained after filtering, where 147 sequences were identified with PhyloNorway, 191 with ArctBorBryo and 308 with EMBL (Supplementary Tables S2-3). Among sequences with less than 10 repeats across all samples, we discarded 64 that had <100% match to the local reference libraries. We assume these mainly represent homopolymer variants of more common sequences, PCR and sequencing artefacts, but it also excluded some taxa that were rare in our dataset (e.g., *Anthyrium*). The remaining 260 sequences were manually cross-checked for consistency. Sequences that were assigned to the same taxon and co-occurring in samples, were merged. Also, 402,835 reads of 40 sequences of known food contaminants or those that were biogeographically unlikely were omitted. We removed four samples from Torfdalsvatn (two of them from the Saksunarvatn tephra layer) and two samples from Nykurvatn due to low quality scores (Supplementary Table S4). This resulted in a final data set of 15,966,705 reads of 191 taxa (146 vascular plants and 45 bryophytes).

### 4.4 Vegetation at Torfdalsvatn

At Torfdalsvatn, we identified 60 taxa of forbs, 31 bryophytes, 23 graminoids, 11 woody, 16 aquatic macrophytes, 4 ferns, 3 horsetails, 1 club moss and 3 taxa of algae, in total 152 taxa (Fig. 5, Supplementary Table S4). The CONISS analyses identified 5 zones (TDV-DNA1 to TDV-DNA5) with breakpoints at approximate 10,100 cal. yr BP, 6700 cal. yr BP, 2400 cal. yr BP and 950 cal. yr BP.

**Fig. 5.**
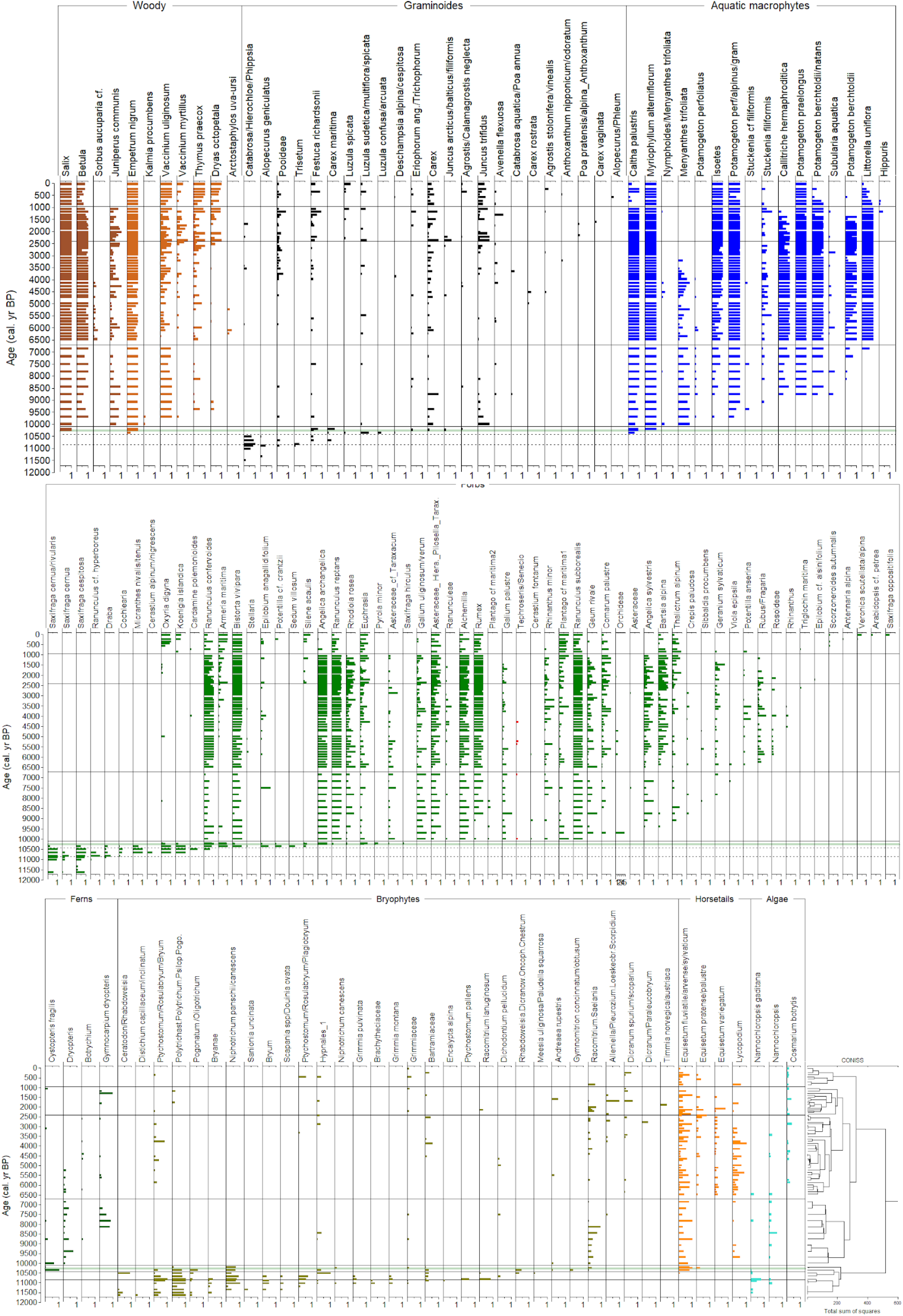
Taxa identified in the *sed*aDNA metabarcoding analyses sorted according to plant functional group (colour bars) and arrival time for Torfdalsvatn, Iceland. All data shown as occurrence in weighted proportion of 8 PCR repeats (see methods). Assumed naturalised non-native taxa are in red and with a star after the name. Zonation according to CONISS analyses are shown with black lines, lithological units with black broken lines and the Saksunarvatn tephra in grey line. Note that some taxa names are shortened. See Supplementary Table S4 for full taxa list.

The oldest CONISS zone (TDV-DNA1, 11,600-10,100 cal. yr BP) included the grey clay, the brown-grey clay, and the start of the light brown-olive gyttja (Fig. 4A). The grey clay was characterised by high arctic forbs (such as *Saxifraga* spp., *Ranunculus, Draba*), grasses (most likely dominated by *Phippisa algida*, but also *Trisetum* and *Alopecurus*) and diverse bryophytes (19 taxa, Fig. 5), with the bryophytes being most abundant (Fig. 6). During this period, the highest proportion of unidentified reads occurs, potentially representing algae which are generally poorly represented in reference libraries. This is also the period with the highest abundance of graminoids in this record, which constitutes up to 34% of the terrestrial plant reads (Fig. 6C, Supplementary data S5). Note however that graminoids may be underestimated with our DNA protocol (Alsos et al., 2018). The diversity increased in the brown-grey clay (from *c*. 10,850 cal. yr BP) with more high arctic forbs like *Oxyria digyna, Koenigia islandica, Cerastium, Micranthes* and *Cardamine polemoioides,* the first appearance of sedge (*Carex maritima*) and seven more bryophytes (Fig. 5). After that, the forbs, mainly *Saxifraga* spp., almost completely replaced the bryophytes for a short period (Fig. 6). Only after the transition to the light brown-olive gyttja (*c*. 10,400 cal. yr BP), the first woody species appear with the dwarf shrub *Empetrum nigrum* and the willow *Salix*, which most likely represent *Salix arctica* or *S. herbacea*. At the same time, 15 new forbs, eight new graminoids (including rushes such as *Luzula* spp.) and three new bryophytes appear. In addition, the first aquatic macrophyte (*Myriophyllum alterniflorum*), fern (*Cystopteris fragilis*), algae (*Nannochloropsis gaditana*), and horsetails (*Equisetum* spp.) appear (Fig. 5).

**Fig. 6.**
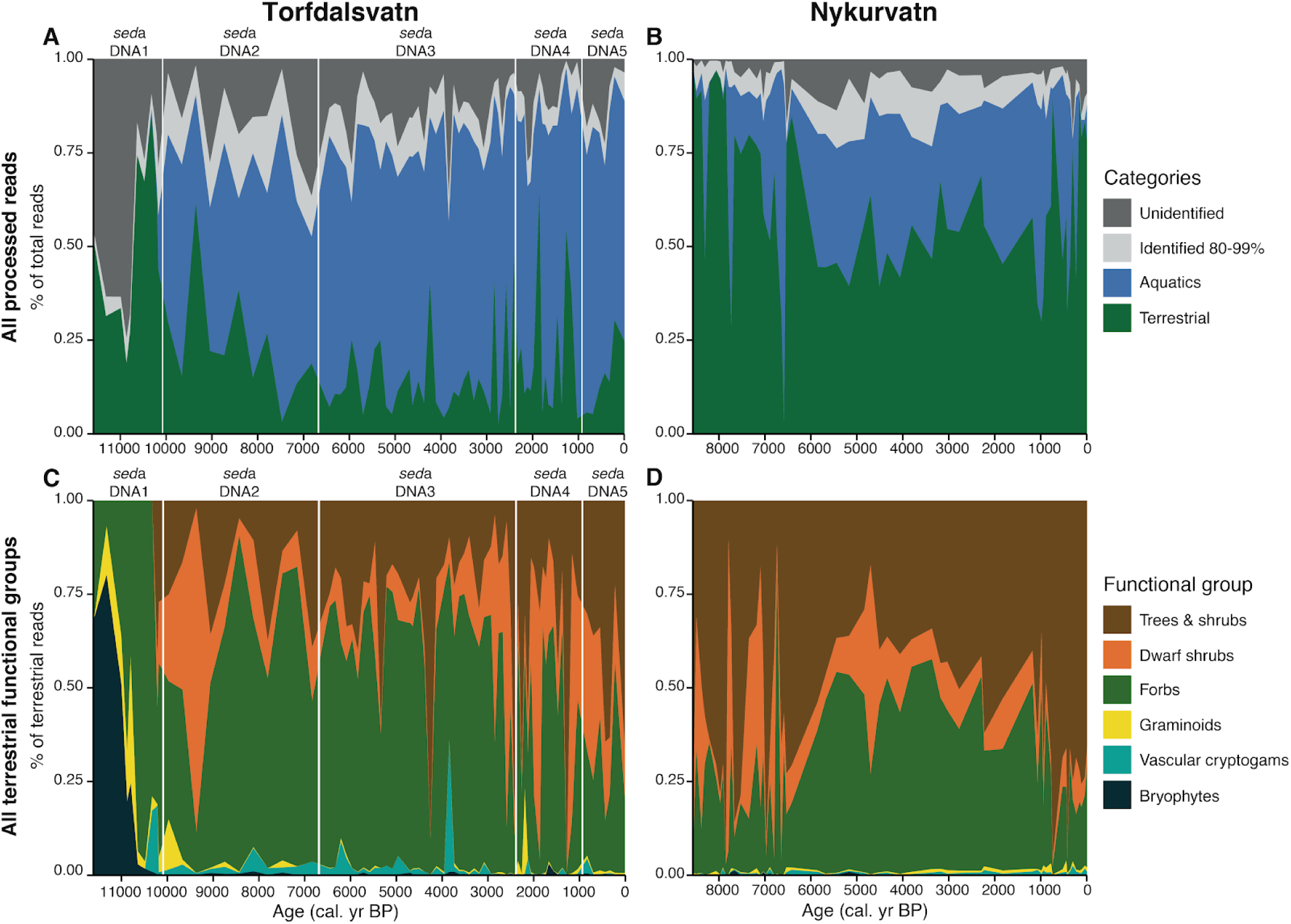
Vegetation development shown as proportion of reads for each functional group. The proportion of terrestrial and aquatic reads is shown for Torfdalsvatn (A) and Nykurvatn (B), along with the proportion of unidentified and partially identified reads. The proportion of reads for terrestrial functional groups is shown for Torfdalsvatn (C) and Nykurvatn (D). For Nykurvatn, the bryophytes, graminoids and cryptogams have average read proportions <0.01 and thus are not clearly visible. Vertical lines in A and C denote the *sed*aDNA zones according to CONISS analyses.

No further change in lithology occurs in the other CONISS zones. Zone TDV-DNA2 (10,100-6700 cal. yr BP) was characterised by a massive immigration of species, with the arrival of *Betula* (either *B. nana* or *B. pubescens*), *Sorbus aucuparia, Juniperus communis* and the dwarf shrubs *Kalmia procumbens, Vaccinium uliginosum, V. myrtillus, Thymus praecox* and *Dryas octopetala* (Fig. 5). In total, 23 new taxa of forbs arrived, the majority of them appearing in most samples onwards, for example *Galium,* Asteraceae, *Alchemilla, Rumex* and *Plantago*, but also some rare forbs such as Orchidaceae. Also, a few new graminoids, ferns, algae, horsetails and one club moss arrive (Fig. 5). This zone is when most aquatic macrophytes arrive, making up >50% of the reads in most samples for the rest of the lake record (Fig. 6A). In addition to the massive immigration, this zone is characterised by the disappearance of many of the early high arctic forbs, *Phippsia*, and bryophytes.

The transition to zone TDV-DNA3 (6700-2400 cal. yr BP) is characterised by arrivals of a few new species. Rowan (*Sorbus aucuparia*) appears in almost every sample in the first two thirds of the zone but then it disappears *c.* 4000 cal. yr BP. The change is most clear in the aquatics, with the aquatic macrophyte *Littorella uniflora* and the algae *Cosmarium botrytis* appearing. Also, a few members of the rose family (*Potentilla anserina, Rubus/Fragaria*) appear. There is a sudden peak in the proportion of trees and shrubs at 4300 cal. yr BP (Fig. 6C), mainly caused by 40% *Juniperus communis* in a single sample (Supplementary Table S5); this may be due to occurrence of macrofossils in that sample rather than a change in the vegetation. There is no disappearance of taxa, but the fern *Gymnocarpium dryopteris* becomes less common.

TDV-DNA4 (2400-950 cal. yr BP) is poorly distinguished from the previous zone and mainly represents a change in abundance of some taxa, such as an increase in *Juniperus communis, Vaccinium myrtillus, Thymus praecox, Dryas octopetala, Festuca richardsonii, Juncus trifidus,* and *Thalictrum alpinum* (Supplementary Table S5). Only scattered records of one new aquatic macrophyte (*Hippuris vulgaris*), two graminoids, one bryophyte and one forb are observed (Fig. 5). However, some of the early high arctic taxa, such as *Silene acaulis* and *Oxyria digyna,* reappear. A reduction in a few taxa, such as *Equisetum variegatum* and *Lycopodium*, is also observed.

The uppermost zone, TDV-DNA5, starts at ∼950 cal. yr BP (AD 1000, taken as midpoint between sample TDV.79 at median age 1055 cal. yr BP and TDV.80 at 846 cal. yr BP). According to the CONISS analyses, the change at 950 cal. yr BP represents the most pronounced change since the onset of the Holocene warming around 10,100 cal. yr BP (Fig. 5). Notably, *Juniperus communis* suddenly disappears after being found in almost every sample from 10,000 cal yr BP to 1055 cal. yr BP (AD 895). A whole range of forbs disappears or reduces in abundance after being common for nearly 9000 years: *Ranunculus confervoides, R. reptans, Angelica archangelica, A. sylvestris, Rhodiola rosea, Galium, Rhinanthus minor and Geum rivale.* In contrast, two of the high arctic forbs, *Koenigia islandica* and *Oxyria digyna*, become common again, together with arrivals of some new arctic-alpine forbs, e.g., *Saxifraga oppositifolia, Antennaria alpina*, and *Scorzoneroides autumnalis*. *Plantago maritima* increases in abundance. Notably, a sequence identified as potential archaeophyte (*Phleum pratense* and/or *Alopecurus pratensis*) appears. Three aquatic macrophytes, *Caltha palustris, Callitriche hermaphroditica* and *Potamogeton berchtoldii,* disappear, indicating a shift in the lake water conditions. The proportion of trees and shrubs increases towards the end of this zone, mainly due to an increase in *Salix* (Figs 5-6).

### 4.5 Vegetation at Nykurvatn

We identified 66 taxa of forbs, 38 bryophytes, 27 graminoids, 15 aquatic macrophytes, 14 woody, 5 ferns, 3 horsetails, 1 club moss and 3 taxa of algae, in total 172 taxa (Fig. 7, Supplementary Table S4). CONISS analyses did not identify any significant zonation in the DNA data. This is in accordance with the uniform lithology (Fig. 4B). All the dominant taxa occurred already in the oldest few samples. Notably, the record starts at 8600 cal. yr BP, which is ∼1500 years after the massive immigration observed at Torfdalsvatn. *Salix* was found in all samples, and *Betula* in all except two, both of which had generally low DNA quality scores. The non-native tall shrub/tree *Alnus* appeared in two samples (5200 and 200 cal. yr BP), whereas the native *Sobus aucuparia* only was found in one sample from 1200 cal. yr BP. *Juniperus communis* was common until 7000 cal. yr BP, after which it only had scattered occurrences, and it was not found in samples from the last 1000 years. Also, some arctic-alpines were most abundant in the early part and then again in the last 1000 years (e.g., *Euphrasia*, *Galium*, *Silene acaulis, Oxyria digyna, Antennaria alpina* and *Pedicularis flammea*). There is a change in aquatic macrophytes, with the more nutrient demanding species *Potamogeton berchtoldii* and *Caltha palustris* only being common from 7600-3300 cal. yr BP. In contrast to Torfdalsvatn, bryophytes were common throughout the core. One taxa classified as naturalised non-native taxon (Wąsowicz, 2020), *Lamium album* (four records 8100-2300 cal. yr BP), was also observed.

**Fig. 7.**
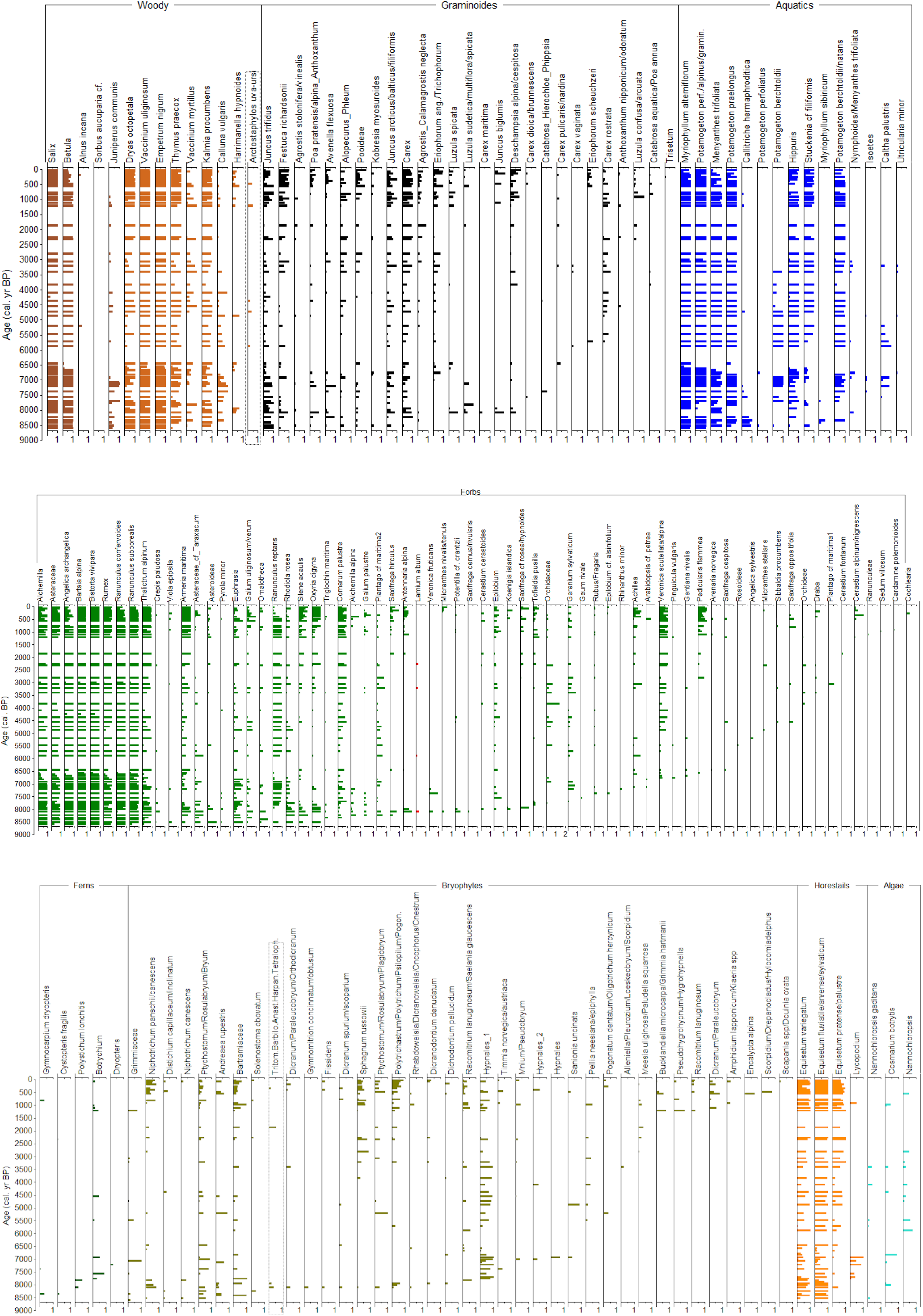
Taxa identified in the *sed*aDNA metabarcoding analyses sorted according to plant functional group (colour bars) and arrival time for Nykurvatn. All data shown as occurrence in weighted proportion of 8 PCR repeats (see methods). One assumed naturalised non-native taxon is in red and with a star after the name. Note that some taxa names are shortened. See Supplementary Table S4 for full taxa list.

### 4.6 Richness and accumulation of species pool

The earliest recorded arrival date of each species is given in Supplementary Table S6. At both sites, there is a rapid increase of species in the early part of the cores. At Torfdalsvatn, there is a steep increase in species richness of bryophytes prior to the increase in forbs, whereas the opposite is observed for Nykurvatn (Fig. 8). At both sites, the richness of trees and shrubs, dwarf shrubs, vascular cryptogams, and aquatic macrophytes levelled off after the first wave of colonisation. The number of forb taxa continued to increase at both sites. The apparent levelling off during some shorter periods at Nykurvatn (e.g., ∼6000, ∼4000, 2500 and 1500 cal. yr BP) may represent sampling artefacts as it coincides with periods with less dense sampling interval (Fig. 7). In contrast, the levelling off at 5900-700 cal. yr BP at Torfdalsvatn is in a period with dense sampling (Fig. 5), so this is assumed to represent a real pattern. Note that the number of forbs increases again at Torfdalsvatn from around 600 cal. yr BP (1350 AD). The number of bryophytes levels off at 10,300 cal. yr BP at Torfdalsvatn, with only four additional bryophyte taxa thereafter. At Nykurvatn, the number of bryophytes continues to increase with a pronounced increase from 1200 cal. yr BP (750 AD). On the contrary, the number of graminoids levels off at 3800 cal. yr BP at Nykurvatn, whereas at Torfdalsvatn graminoids increases up to 8800 cal. yr BP and then again from 5000 cal. yr BP.

**Fig. 8.**
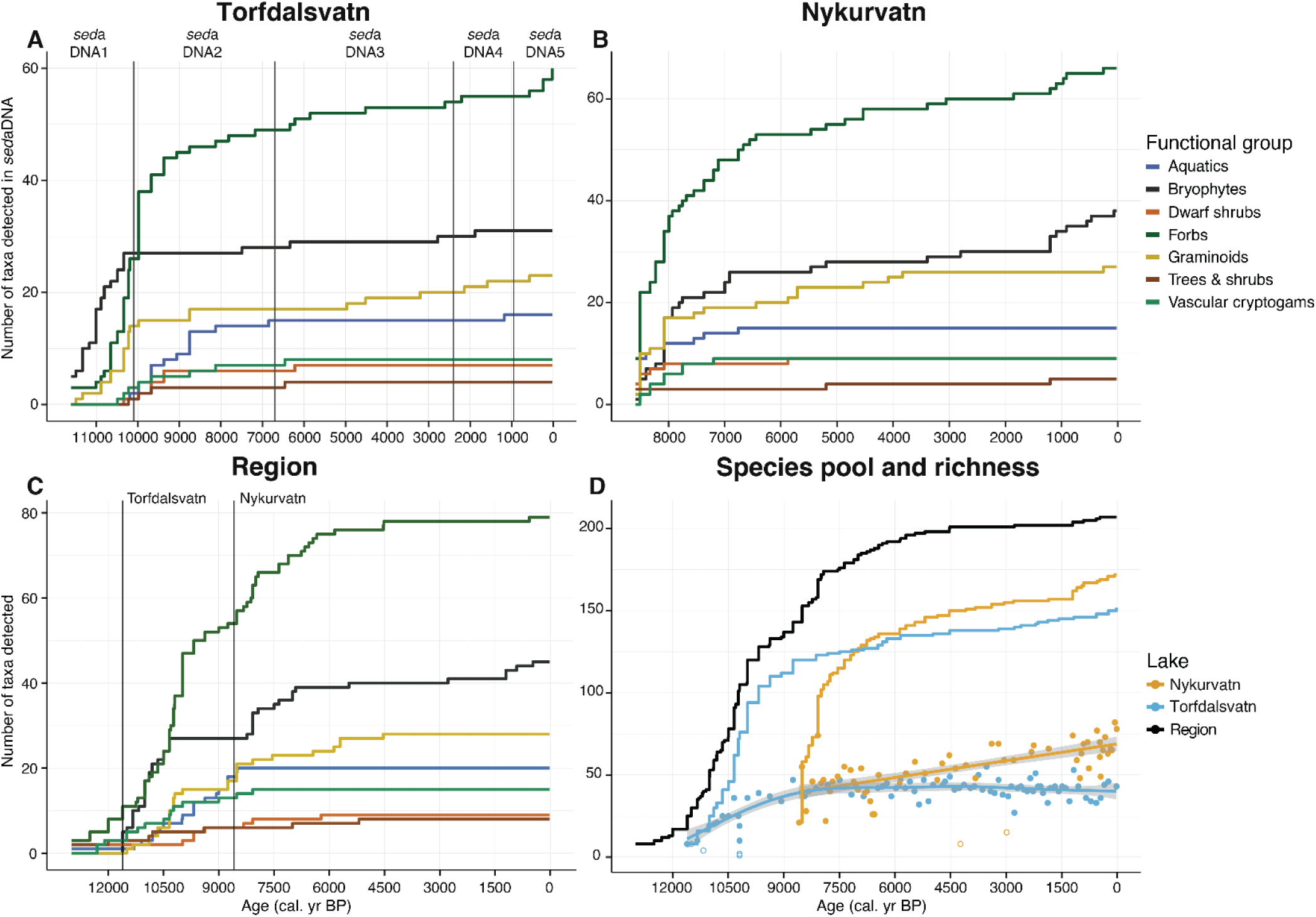
Development in species richness, local and regional species pools in Iceland. Species pool (defined as cumulative number of taxa) per functional type for Torfdalsvatn (A), Nykurvatn (B), and both sites combined with the macrofossil and pollen data from Alsos et al., 2016 (C). Vertical lines in A denote the *sed*aDNA zones according to CONISS analyses, while the vertical lines in C indicate the start of the Torfdalsvatn and Nykurvatn cores. (D) Accumulated detected regional species pool as well as number of taxa detected per sample along with the 95% confidence interval (grey shading) of the fitted line. Open circles represent excluded samples, which were not used for the richness curves (Supplementary Table S6). The regional species pool is the data from both sites combined, along with the macrofossil and pollen data from (Alsos et al., 2016).

When data from both sites as well as the review study are aggregated, only 1-2 new taxa of vascular cryptogams, trees and shrubs, dwarf shrubs or aquatic macrophytes are observed after 8000 cal. yr BP (Fig. 8C). No new graminoids and only one new forb arrive after 4500 cal. yr BP, whereas bryophytes increase in the last 1000 years.

The average raw richness was 13.92 for Torfdalsvatn and 19.19 for Nykurvatn. The richness per sample levelled off from around 8000 cal. yr BP at Torfdalsvatn, whereas it showed an almost linear increase of 3.5 taxa per millennium for Nykurvatn (Fig. 8D). The species pool at Torfdalsvatn showed a steep increase from 11,600 to ∼9000 cal. yr BP, and a continuous increase after that (Fig. 8D). The initial increase at Nykurvatn was even steeper, and continues to increase until present. The detected regional species pool based on both lakes and review of earlier records shows a steep increase until around 8000 cal. yr BP, when >80% of the total taxa had arrived, and hardly any new taxa after 4500 cal. yr BP (Fig. 8D).

## 5. Discussion

Our records characterising Early Holocene vegetation and plant communities reflect summer temperature conditions in accordance with previous Holocene Thermal Maximum reconstructions (Caseldine et al., 2006; Eddudóttir et al., 2015; Hallsdóttir and Caseldine, 2005; Harning et al., 2018). Fig. 9 summarises key vegetation data from each site compared to sea-ice proxy IP_25_ data from the North Iceland shelf (Cabedo-Sanz et al., 2016; Xiao et al., 2017) and reconstructed mean summer air temperature anomalies from Northwest Iceland (Harning et al., 2020). The largest changes in taxonomic composition and dominance were observed at Torfdalsvatn at the onset of the Holocene warming around 10,100 cal. yr BP, coinciding with peak summer insolation (Berger and Loutre, 1991), and at Landnám about 1000 years ago. The Nykurvatn record starts at 8600 cal. yr BP, and exhibits the largest shift following Landnám, although without statistical significance. Furthermore, minor changes were observed in both records, mainly in the aquatic macrophytes. While the taxonomic composition was rather stable from 10,100-1000 cal. yr BP, the proportion of different plant functional groups fluctuated. The majority of taxa immigrated before 8000 cal. yr BP, thus prior to the Holocene Thermal Maximum (Geirsdóttir et al., 2020). The peak immigration was around 10,000 cal. yr BP, when (Fig. 9), when occasional sea ice connected Iceland to northern Scandinavia and Russia (Alsos et al., 2016).

**Fig. 9.**
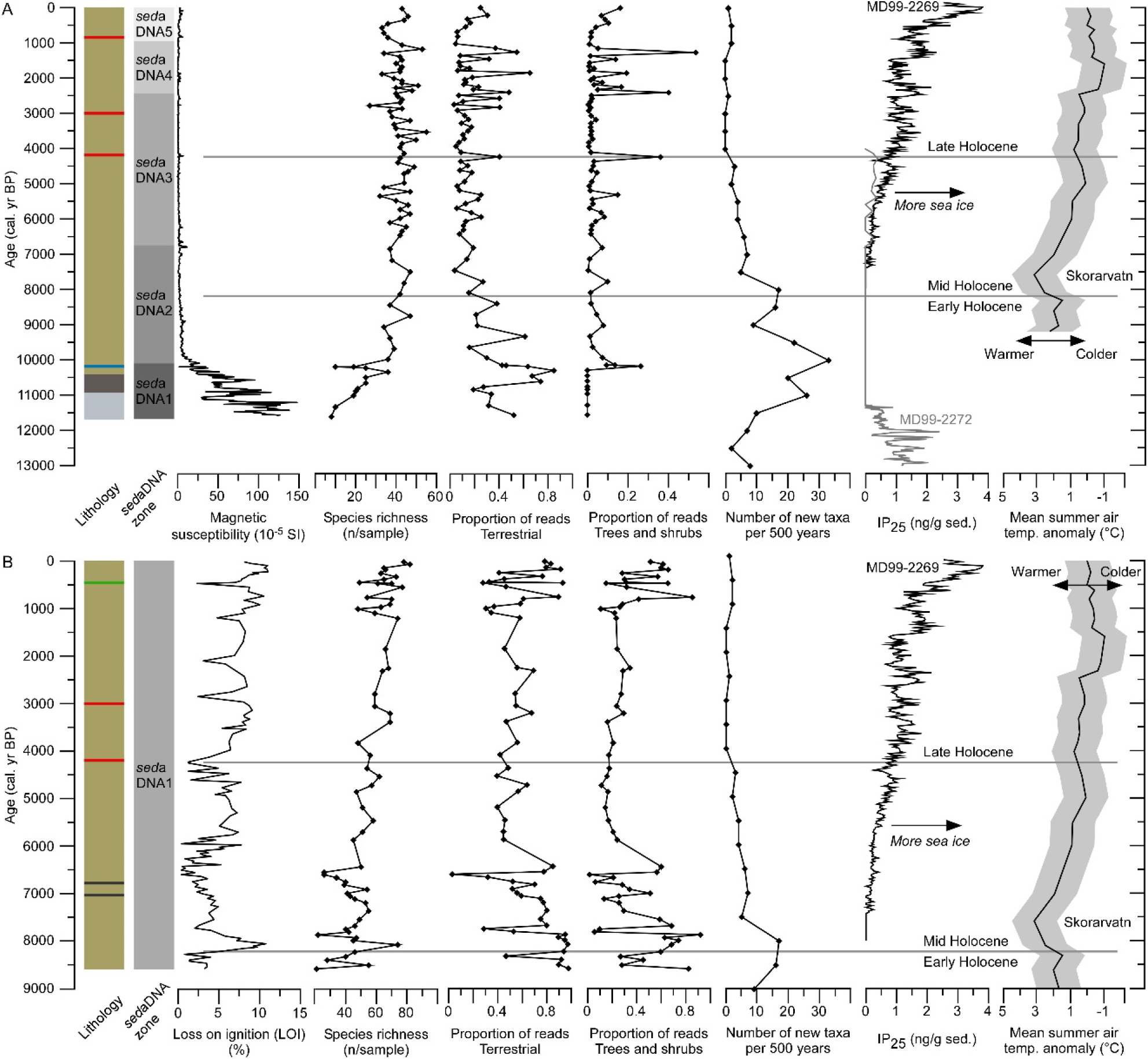
Lithology, *sed*aDNA zones from the CONISS analyses, magnetic susceptibility (MS, Torfdalsvatn only), loss on ignition (LOI, Nykurvatn only), species richness, proportion of reads for terrestrial plants, proportion of reads for trees and shrubs, and number of new taxa recorded per 500 years (new taxa from both sites and Alsos et al., 2016) compared to regional paleoclimate records. The IP_25_ biomarker record is a proxy for sea ice from two marine sediment cores at the North Iceland shelf (MD99-2269, Cabedo-Sanz et al., 2016; MD99-2272, Xiao et al., 2017, note that even 11,000-7000 cal. yr BP, some sea ice was present NE of Iceland occasionally connecting Iceland to N Scandianvia/Russia, Alsos et al., 2016). The mean summer air temperature anomaly is based on the biomarker brGDGT from Skorarvatn, Northwest Iceland (Harning et al., 2020). Grey shading indicates the propagated standard error. The key to the lithology is given in Fig. 4. Results from Torfdalsvatn (A) and Nykurvatn (B).

### 5.1 Possible glacial survival of high arctic species

The assemblage of high arctic taxa found in the oldest samples at Torfdalsvatn, suggests that these might have survived the LGM in Iceland. However, as we have no sediments older than *c.* 11,600 cal. yr BP, we have no direct proof of glacial survival. Glacial survival of high arctic forbs, graminoids and bryophytes is less controversial and has also been found in a LGM refugium in northern Norway (Alsos et al., 2020). In our current study, possible glacial survival taxa based on their occurrence in the three oldest samples are *Saxifraga cespitosa, Saxifraga cernua* and/or *rivularis* and *Phippsia algida*. For *S. cernua,* a considerable intermingling in the amphi-Atlantic region was inferred, whereas no genetic variation was detected within the region in *P. algida* (Aares et al., 2000; Brochmann et al., 2003; Bronken, 2001). We are not aware of any phylogeographical studies of *S. cespitosa* that include Iceland, but a study of populations in Svalbard and Norway concludes high dispersal potential (Brochmann et al., 2003; Tollefsrud et al., 1998). Notably, potential glacial survival has been suggested for *S. rivularis* in Svalbard, but not in Iceland, based on studies of AFLP fingerprinting (Westergaard et al., 2010). Thus, our new data neither prove nor disprove glacial survival.

### 5.2 Colonisation of plants

The initial steep increase in accumulated species richness at both sites represents an overestimation of colonisation at that time. For Torfdalsvatn, we know that many of the taxa were present earlier (Rundgren, 1998, 1995), and for Nykurvatn, our record does not include the earliest vegetation in the region. Thus, numerous species arrived prior to their being detected in our *sed*aDNA records (Figs 8-9). Also, it takes time from the first arrival of a propagule to become spread around Iceland. Therefore, it is likely that the massive immigration mainly took place prior to 10,100 cal. yr BP. Sea ice between Iceland and North Norway was common to dense prior to 11,000 cal. yr BP (Alsos et al., 2016). In addition, catastrophic draining of the Baltic Ice Lake around 11,700 cal. yr BP caused meltwater and potentially ice loaded with sediments to drift into the Nordic Seas (Björck, 1995; Nesje et al., 2004). Both sea ice and drift ice have already been identified as a major dispersal vector for beetles to Iceland (Panagiotakopulu, 2014). After that, sea ice was only found occasionally around Iceland 11,000-8000 cal. yr BP (Alsos et al., 2016), and may have been completely absent on the North Iceland shelf 11,700-6800 cal. yr BP (Fig. 9, Xiao et al., 2017). This direct evidence of a massive immigration prior to 10,100 cal. yr BP. supports that sea ice may have acted as an important dispersal vector either by direct transport of diaspores or by a combination of wind and sea ice (Alsos et al., 2016; Panagiotakopulu, 2014), although other dispersal vectors as birds, wind and sea current also may have contributed (Alsos et al., 2015).

From around 8000 cal. yr BP, hardly any taxa of trees, shrubs, dwarf shrubs, aquatic macrophytes or cryptogams were detected in the lake records. These are functional groups that have rather few taxa in Iceland (Wąsowicz, 2020). There are, however, more taxa of these groups in the assumed main source regions Scandinavia and the British Isles (Elven, 2005; Stace, 1997). As aquatic macrophytes are well dispersed by birds and appeared within 1-3 decades after warming in western Norway (Birks, 2000), they may not be limited by availability of sea ice. They may, however, be limited by nutrients and light availability or other factors related to habitat diversity in Iceland. Trees, shrubs, dwarf shrubs, and cryptogams on the other hand, might have been more limited by lack of sea ice in the Middle Holocene.

In contrast, new taxa of forbs, graminoids, and bryophytes continued to be detected until 4500 cal. yr BP (bryophytes until present). Forb is the functional group with the highest number of taxa in total (Fig. 8). The majority of forbs rely on insects for pollination. Thus, late colonisation of forbs may be due to pollinator limitation. The arrival time for most pollinators is not known (Panagiotakopulu, 2014). However, we note that most forb taxa and most graminoids that are only recorded after 4500 cal. yr BP are found scattered in a few samples and only in a few of the PCR repeats (Figs 5 and 8). As the ability to detect rare taxa in *sed*aDNA is limited (Alsos et al., 2018), they may represent scattered records of taxa that were also present earlier. Indeed, *Saxifraga oppositifolia* was recorded on Svalbard and Greenland 11,000-10,000 cal. yr BP (Bennike et al., 1999; Bennike and Hedenäs, 1995), so it is not unlikely that it was also present in Iceland at that time. The late arriving bryophytes however, were detected in many samples and often all eight PCR repeats, suggesting that these taxa are less likely to have been overlooked in earlier samples. Bryophytes have small spores that in general are widely dispersed. Furthermore, they are important pioneers on volcanic substrate, producing organic material for secondary colonisers (Ingimundardóttir et al., 2014). While late arrival of some species cannot be excluded, the cooler conditions during the Little Ice Age (AD 1250/1500-1900, Mann, 2002) may have made bryophytes more abundant and thus more likely to be detected.

The overall accumulation of plant taxa in Iceland is greatest during the Late Glacial and Early Holocene (>80% before 8000 cal. yr BP). Furthermore, these records from Iceland exhibit an earlier accumulation compared to an accumulation curve for plant taxa based on *sed*aDNA analyses of ten lakes in northern Fennoscandia (⅔ before 8000 cal. yr BP, Rijal et al., 2020). This could be related either to species source pools or dispersal vectors. The flora in Iceland’s main source regions, Scandinavia (Elven, 2005) and the British Isles (Stace, 1997), are about four times as large as the flora of Iceland (Alsos et al., 2015; Wąsowicz, 2020). While some of these species require warmer conditions than currently found in Iceland, it is clear that there is a large species pool that never made it across the ocean. Thus, we do not think that the size of the source pool *sensu* (Harrison, 2020) has limited colonisation of plants in Iceland. As our *sed*aDNA record is based on only two lakes, both lacking the basal sediments, the difference between Iceland and northern Fennoscandia is likely underestimated. The accelerated accumulation of taxa in Iceland compared to northern Fennoscandia may relate to the disappearance of the main dispersal vector (sea ice) during the Middle Holocene, and therefore we did not observe the same dispersal lag as in northern Fenoscandia (Rijal et al., 2020).

Ultimately, if sea ice has been an important natural vector for plant dispersal to Iceland as our data suggest, we may not expect a wave of immigration of new taxa to Iceland due to the decreased ice cover following ongoing climate change. Furthermore, most modern sea ice originates from the Arctic Ocean, and is transported south through the Denmark Strait by the East Greenland Current. Anthropochore introduction, however, is assumed to increase unless measures are taken (Wasowicz et al., 2013).

### 5.3 Early Holocene vegetation development

The Saksunarvatn tephra layer provides a firm tie point between our TDV core and cores from Torfdalsvatn previously investigated by Rundgren (1995) and Florian (2016), and magnetic susceptibility provided additional alignment to the core by Florian (2016). As in the Rundgren (1995) study, the Saksunarvatn tephra overlaid a layer of clayey gyttja, which again overlays a layer of clay. We did not observe the Vedde tephra layer (12,100 cal. yr BP, Rasmussen et al., 2006), thus TDV only extends down to somewhere in the clay layer (pollen zone T4-T5) of Rundgren (1995).

The lowest part of our core (TDV-DNA1) is dominated by bryophytes with some *Saxifraga*, which aligns with the T4 zone of low pollen concentrations detected by Rundgren (1995). This is also the period with the highest proportion of unidentified sequences (Fig. 6). As the majority of bryophytes and vascular plants known from Iceland are covered in our reference library, the unidentified sequences likely reflect algae, which have limited reference material available for the marker used. This aligns with assumed high autochthonous production of algae based on algal pigments and a low ratio of total organic carbon to biogenic silica (Florian, 2016).

In our core, the bryophytes are rapidly replaced by high proportions of *Saxifraga* species, while graminoids remain common and peak around 10,800 cal. yr BP. This likely corresponds to the increase in grass pollen exhibited in pollen zone T5 of Rundgren (1995). Thereafter, both *sed*aDNA and pollen (T6) show an overall high number of taxa; such as *Salix, Empetrum nigrum, Oxyria digyna, Koenigia islandica, Ranunculus,* Caryophyllaceae (identified to *Cerastium alpinum/nigrescens* and *Silene acualis* in *sed*aDNA), and *Hornungia* type pollen (identified to *Cardamine polemonioides* in *sed*aDNA)(Rundgren, 1998, 1995). This is a period characterised by increasing ratios of carbon to nitrogen and total organic carbon to biogenic silica (Florian, 2016), suggesting enhanced terrestrial production.

Our TDV-DNA2 (10,100-6700 cal. yr BP) is characterised by the appearance of *Juniperus communis*, which is assumed to correspond to pollen zone T8. In contrast to the previous record (Rundgren, 1998), we observed a massive immigration of new taxa at this transition, especially of forbs which are generally poorly represented in pollen. The transition to pollen zone T9 at around 8700 cal. yr BP is mainly quantitative, and could not easily be aligned with any change in the DNA data (falls within TDV-DNA2). Note that the two dates Rundgren has from T9 are reversed (7940+/-130 and 7860+/-120 BP). Also, there is a 4000 year hiatus in the record of Rundgren (1998), so his analyses stop here. Soon after, the first aquatic macrophytes appear in our DNA record, and they become the most dominant group in terms of DNA reads. This aligns with the increase in δ^13^C just after deposition of the Saksunarvatn tephra layer, supporting an increase in organic production by aquatic macrophytes (Florian, 2016) consistent with a clearer water column following the reduced minerogenic influx also shown here by the MS profile (Fig 9A). The δ^13^C remains higher than -20°C until the last millennium (Florian, 2016), which agrees with the high proportion of DNA reads identified as aquatic macrophytes. The high proportion of aquatic macrophytes may also explain the lower richness detected at Torfdalsvatn compared to Nykurvatn, as the richness of terrestrial taxa may be underestimated when >50% of the DNA reads are originating from aquatic taxa (Rijal et al., 2020).

### 5.4. Vegetation development during Middle and Late Holocene

At Nykurvatn, no significant changes in vegetation composition were found for the entire period of 8600 years, suggesting a very stable flora. While we cannot distinguish *Betula nana* from *B. pubescens* based on the DNA region studied, the lack of other forest taxa (e.g., few ferns) and the scattered occurrence of *Juniperus communis* (<8 PCR repeats except around 8000-7000 cal. yr BP) suggest that forest never developed at this site. Thus, this was therefore not affected by oscillations in *Juniperus communis* and *Betula* cover. This interpretation is supported by pollen studies from the northwestern highland (416 m a.s.l), where sensitivity to climate and tephra deposition is demonstrated by floral communities in which the thermophilous (Iceland) *Betula pubescens* is prominent (Eddudóttir et al., 2016, 2017). The variations in LOI and Ti/Sum are likely caused by the numerous silt-sand sized tephra beds as well as input of minerogenic material from runoff and aeolian activity, rather than major environmental change within this period.

The occurrence of the non-native tall shrub/tree *Alnus* at 5200 and 200 cal. yr BP is somewhat unexpected, especially as these do not fall within a warm period (Geirsdóttir et al., 2020). It may be difficult to distinguish small amounts of *sed*aDNA from background contamination (Alsos et al., 2020). However, we have never recorded *Alnus* as contamination in our laboratorium so far (Rijal et al., 2020). Also, the scattered occurrence of *Alnus* in some pollen records (Verhoeven and Louwye, 2015) suggests that it might have occurred in Iceland in the past.

At Torfdalsvatn, the transition to zone TDV-DNA3 (6700-2400 cal. yr BP) represents a minor change in the terrestrial vegetation, but rowan appears from 6500-4100 cal. yr BP. Rowan also appears in pollen diagrams from northern and southern Iceland from ∼6000 and 4500 cal. yr BP (Eddudóttir et al., 2015; Hallsdóttir, 1995; Hallsdóttir and Caseldine, 2005). This tree species has a slightly more southern distribution than *Betula pubescens.* It is believed that its distribution is not limited by summer temperature *per se*, but rather by a combination of poor drought tolerance, adaptation to short growing seasons and cold requirement for bud burst (Raspe et al., 2000). This suggests that the winters were colder during this period, which agrees with the re-occurrence of sea ice at the North Iceland shelf after *c*. 6800 cal. yr BP as indicated by IP_25_ biomarker data (Xiao et al., 2017), and by the presence of quartz in marine sediment cores after *c*. 7000 cal. yr BP (Andrews et al., 2009). The disappearance of rowan around 4100 cal. yr BP is during a period with widespread ice cap expansion, decreasing biogenic silica and increased landscape instability (Geirsdóttir et al., 2020).

There was a clear change in the aquatic flora of Torfdalsvatn with the aquatic macrophyte *Littorella uniflora* appearing from 6800 cal. yr BP. This is approximately the same time as δ^15^N increases, δ^13^C decreases, and there is a shift in algal pigments (Florian, 2016), possibly related to lacustrine environmental changes. *Littorella uniflora* is rare in Iceland and not detected in Skagi today (Kristinsson, 2010) although its occurrence in the top samples suggests it is overlooked.

The change in abundance of some taxa at the transition to TDV-DNA4 (2400 cal. yr BP) with scattered reappearance of some of the early high arctic taxa, such as *Silene acaulis* and *Oxyria digyna,* may be a result of the stepwise cooling that started in Iceland around 5000 cal. yr BP, with increased cooling around 2400 cal. yr BP (Geirsdóttir et al., 2020, 2019; Harning et al., 2020). A similar change was also observed at Nykurvatn, where *Oxyria digyna* and *Silene acaualis* increased from ∼2000 cal. yr BP, and several arctic-alpines re-appeared.

### 5.5 Landnám, Medieval Climate Optimum and Little Ice Age

The clearest indication we observed of human impact was the sudden disappearance of *Juniperus communis* 1055 cal. yr BP (895 AD) at Torfdalsvatn and 910 cal. yr BP (1040 AD) at Nykurvatn. Some grazing may favour this species and it is therefore commonly taken as a grazing indicator, especially of dry pastures (Vorren, 1986), although grazing reduces seedling survival (Thomas et al., 2007). The disappearance from the DNA record suggests that in addition to grazing, it was cut. This is a species that was used as fuel, fodder for sheep and as raw material for ropes and tools (Larsen and Others, 1990). A reduction in *Juniperus communis* has also been observed in pollen records from *c*. 1000 cal. yr BP (Eddudóttir et al., 2020, 2016). Additional indicators of Landnám were the increase in a few grass taxa at Nykurvatn, as well as the reduction in *A. sylvestris* and disappearance of *Angelica archangelica* at Torfdalsvatn. These are taxa that are known to be grazing sensitive and also decrease or disappear in pollen records (Eddudóttir et al., 2020; Erlendsson, 2007). The latter was also observed at Nykurvatn, but continued to grow there during the last 1000 years, probably because this is a higher altitude site less influenced by human land use. While the onset of the Medieval Climate Optimum coincides with Landnám, we did not see any strong indication of more thermophilic flora. Thus, the change at around 950 cal. yr BP at both sites (but only significant at Torfdalsvatn), is interpreted as mainly due to human impact.

The increase of high arctic taxa such as *Oxyria digyna, Koenigia islandica, Epilobium anagallidifolium, Silene acaulis,* and *Dryas octopetala* at Torfdalsvatn and increase in bryophytes at Nykurvatn occur during the Little Ice Age. Thus, in addition to the impact of human land use, we see a clear effect of climate deterioration. This coincides with low summer air temperatures and increased sea-ice cover on the North Icelandic shelf (Fig. 9; Cabedo-Sanz et al., 2016; Miles et al., 2020).

## 6. Conclusions

By using *sed*aDNA, we obtained detailed records of arrival of species and the vegetation history at one lowland and one higher-altitude lake in Iceland. Our records confirm the presence of a high arctic vegetation prior to peak Holocene warming. Moreover, the high taxonomic resolution of these records allows for the estimation of arrival time. Over 80% of the recorded taxa had arrived by 8000 cal. yr BP, prior to the Holocene Thermal Maximum in Iceland. The main immigration event coincides with a period of more extensive sea ice, supporting the hypothesis that sea ice was an important dispersal vector to Iceland. To further improve estimates of arrival time, more sites should be investigated across the island, focusing especially on records that span the Late Glacial/Holocene transition. We found that the taxonomic composition of the flora was remarkably stable until Landnám, when some land-use intolerant taxa gave way rapidly. While ongoing global warming will allow more species to thrive in the Icelandic environment, natural colonisation may be limited by reduction in sea ice resulting in a shift to anthropochore dominated dispersal.

## Acknowledgements

We would like to thank Lasse Topstad and Luca Elliott for help with DNA extraction, Dilli P. Rijal for help with R script for data analyses, and Ellen Elverland for help with identifying macrofossils. Claus Falkenberg Thomsen kindly assisted during fieldwork at Nykurvatn. Þorsteinn Jónsson and Höskuldur Þorbjarnarson assisted with coring at Torfdalsvatn. Kurt H. Kjær and Marie-Louise Siggaard-Andersen, GLOBE Institute, kindly contributed with ITRAX sediment core scanning. Bioinformatic analyses were performed on resources provided by UNINETT Sigma2 - the National Infrastructure for High Performance Computing and Data Storage in Norway. This project has received funding from the European Research Council (ERC) under the European Union’s Horizon 2020 research and innovation programme (grant agreement No 819192 to Alsos), from the Arctic Research and Studies Program of the Ministries for Foreign Affairs of Norway and Iceland (grant agreement No. 2017-ARS-79772 to Schomacker), the Icelandic Research Fund (grant no. 141842-051; Guðrún Gísladóttir), Landsvirkjun Energy Research Fund and the University of Iceland Research Fund.

## Author contributions

Project idea IGA and AS; acquiring funding IGA, AS, EE, GG, ÍÖB, SB, WRF; coring SB, ÍÖB, EE, GG, SDE, WRF, AS; core sampling EMB, AR; lithology SEK, EMB, AR, AS; identification of plant macrofossils for radiocarbon dating EMB, SEK, AS; analyses of tephra EMB, ERG, WRF; age-depth modelling EMB, SEK, AS; DNA analyses MKFM and IGA; data analyses IGA and YL; manuscript draft IGA, AS, YL and SEK. All co-authors commented on the manuscript.

## Data availability

Raw sequence read data will be uploaded on Dryad (XXXX). The data obtained after filtering are given in Supplementary data Table 4-5.

## Supplementary data

**Supplementary Table S1.** Results from major element analyses on identified tephra layers and standard runs.

**Supplementary Table S2.** The number of unique barcodes and reads present in the data after each bioinformatic step.

**Supplementary Table S3.** Sequence identification with comments.

**Supplementary Table S4.** Final *sed*aDNA dataset for Torfdalsvatn (TDV) and Nykurvatn (NYK) given as the weighted number of PCR repeats.

**Supplementary Table S5.** DNA raw and filtered reads, technical and ecological quality scores and proportion of reads for functional groups and key species for Torfdalsvatn (TDV) and Nykurvatn (NYK).

**Supplementary Table S6.** First arrival of taxa based on the current study and previous review of all macrofossil and pollen studies (Alsos et al., 2016).

**Supplementary Figures S1-S14**

